# The up-regulation of TGF-β1 by miRNA-132-3p/WT1 is involved in inducing leukemia cells to differentiate into macrophages

**DOI:** 10.1101/2024.06.13.598949

**Authors:** Zhimin Wang, Chaozhe Wang, Danfeng Zhang, Xidi Wang, Yunhua Wu, Ruijing Sun, Xiaolin Sun, Qing Li, Kehong Bi, Guosheng Jiang

## Abstract

Although it has been shown that abnormal expression of Wilm’s tumor gene 1 (WT1) is associated with the occurrence of leukemia, the specific mechanism via which it induces leukemia cells to differentiate into macrophages remains poorly understood. Based on the prediction that the microRNA miRNA-132-3p is the miRNA that possibly lies upstream of the WT1 gene, we hypothesized that miRNA-132-3p may participate in the polarization process of macrophages through regulating expression of the WT1 gene. The focus of the present study was therefore to investigate the role of the miRNA-132-3p/WT1 signaling axis in the differentiation of THP-1 leukemia cells into macrophages induced by PMA. The results obtained indicated that, compared with the control group, the proliferation of THP-1 cells was clearly inhibited by PMA, and the cell cycle was arrested at G0/G1 phase, associated with an upregulation of CD11b and CD14 expression. Induced by PMA, the expression level of miRNA-132-3p was increased, WT1 expression was decreased, and the expression level of TGF-β1 was increased. Following transfection with miRNA-132-3p mimics, however, the expression of WT1 in the THP-1 cells was downregulated, with upregulation of the CD11b and CD14 antigens, whereas this downregulation of WT1 mediated by miRNA-132-3p mimics could be reversed by co-transfection with WT1 vector, which was accompanied by downregulation of the CD11b and CD14 antigens. The luciferase activity of the co-transfected miRNA-132-3p mimic + WT1-wild-type (WT) group was found to be statistically significantly lower compared with that of the co-transfected miRNA-132-3p mimic + WT1-mutated (MUT) group. Furthermore, chromatin immunoprecipitation experiments showed that WT1 was able to directly target the promoter of the downstream target gene TGF-β1, which led to the negative modulation of TGF-β1 expression, whereas downregulation of WT1 led to an upregulation of the expression of TGF-β1, which thereby promoted the differentiation of THP-1 cells into macrophages. Taken together, the present study has provided evidence, to the best of the authors’ knowledge for the first time, that the miRNA-132-3p/WT1/TGF-β1 axis is able to regulate the committed differentiation of leukemia cells into macrophages.

## Introduction

Acute myeloid leukemia (AML) is a common malignant hematological tumor originating from promoter cells (1-3). In clinical practice, cytotoxic chemotherapy regimens remain the predominant method for treatment of patients with AML (4). However, resistance to these chemical drugs often develops; moreover, damage frequently occurs to the normal hematopoietic and immune cells, which motivates researchers to find safer and more effective treatment methods. It has been shown that the occurrence of leukemia is to a large extent caused by the differentiation block of pluripotent hematopoietic stem cells at a certain stage during the maturation process, and so patients with leukemia may be treated with inducers of differentiation in order to induce the leukemia cells to differentiate into more mature cells (5,6). For example, all-*trans* retinoic acid (ATRA) has been used to treat patients with acute promyelocytic leukemia (APL) (7), and different leukemia cell lines have been shown to differentiate into mature monocytes, erythrocytes or granulocytes, depending upon their induction by different inducers (8-12). However, in spite of the success of ATRA in inducing APL cells to differentiate into granulocytes, to date, no breakthroughs have been achieved in terms of pathways for the directed differentiation of monocytes or macrophages. As a result, the identification of novel differentiation intervention targets or differentiation-inducing drugs is of crucial importance, and studying the underlying molecular mechanism(s) of abnormal cell differentiation will provide the basis for discovering novel targeted differentiation-inducing drugs.

When confronted with the very complex processes of pathogenesis of leukemia, it is imperative to select suitable molecular targets and relevant signaling pathways. After performing a literature review, our research group found that Wilm’s tumor gene 1 (WT1) is closely associated with the occurrence and development of solid tumors or hematological tumors, suggesting that it may be involved in the committed differentiation process of leukemia cells. WT1 is located on chromosome 11p13, and encodes a DNA-binding protein (13). The carboxyl-terminus of the protein encoded by this gene contains four zinc finger domains, which acts a DNA-binding functional region. It is able to identify and bind to the GC-enriched fragment (CCCCCGC) of the target gene in its promoter region, whereas the amino-terminus is rich in glutamine/proline residues, mainly serving a role in transcriptional regulation (14,15). It has been reported that WT1 is an effective transcription factor that exerts multiple roles in the proliferation, differentiation, development and apoptosis of normal and tumor cells (16,17). To date, studies have shown that WT1, as a tumor gene, is usually upregulated in solid tumors or hematological malignancies (18), and WT1 protein has a crucial role as a regulator of transcription, participating in the proliferation, differentiation and apoptosis of leukemia cells (19,20), as well as having an association with the diagnosis, minimal residual disease (MRD) and prognosis of patients with leukemia (20-22). In terms of the clinical association between WT1 and patients with leukemia, it has been shown that mutations of WT1, and overexpression of the protein, may occur in different subgroups of AML (20). WT1 can be used as a biomarker variously for the diagnosis of AML, monitoring of MRD, clinical management, molecular remission or recurrence detection of the disease (21,22). Furthermore, the high expression status of WT1 provides useful opportunities for the development of vaccines, targeted antibodies or chimeric antigen receptor immunotherapy (23,24). However, the exact role of WT1 in the development of leukemia remains poorly understood; moreover, the complexity associated with this function of WT1 will require a great research effort in the future (25). Even though some research groups have found that the expression of WT1 is downregulated in leukemia cells induced by differentiation inducers (26-28), to the best of the authors’ knowledge, few studies have been published on either the committed differentiation of WT1 in leukemia, or on the committed differentiation of leukemia cells into macrophages.

Numerous studies have shown that oncogenes are often regulated by upstream noncoding RNA or transcriptional regulatory proteins; for example, microRNAs (miRNAs) usually negatively modulate the expression of their target genes, or otherwise regulate important biological processes, such as cell differentiation and apoptosis (29-31). We hypothesized that a certain miRNA may be involved in regulating the expression of WT1. Among these miRNAs, miR-132 is located on chromosome 17, and consists of two homologous miRNAs, namely hsa-miR132-5p and hsa-miRNA-132-3p. Although it has been demonstrated that miR-132 may be a valuable biomarker candidate for the prognosis of various solid tumors (32-41), to date, few studies have been published on its role in leukemia. Preliminary experiments and database analyses have indicated that miRNA-132-3p is associated with the occurrence of leukemia, and it is also a possible upstream miRNA that targets the 3’-untranslated region (3’-UTR) of WT1. However, further in-depth studies are required to delineate how it may regulate WT1, and modulate the differentiation of leukemia cells. In addition, transforming growth factor-β (TGF-β) has also been shown to regulate various cellular activities, including apoptosis, differentiation and angiogenesis, and TGF-β is known to participate in the occurrence of tumors (42,43) and in leukemia (44). Furthermore, TGF-β1 can interact with the receptor of TGFβ1 to participate in cell proliferation, differentiation and apoptosis, or to promote the differentiation of leukemia cells (45). Interestingly, through database analysis and differential gene chip screening, we found that WT1 may be a target gene for hsa-miRNA-132-3p, and that WT1 is associated with downstream TGF-β1 in the pathway.

However, the role of the miRNA-132-3p/WT1/TGF-β1 axis in inducing leukemia cells to differentiate into macrophages has yet to be fully elucidated. Therefore, the aim of the present study was to investigate whether there is a targeted relationship, and any regulatory effects, between miRNA-132-3p and WT1, and subsequently to determine: i) whether WT1 is target-associated with TGF-β1; and ii) whether it has a regulatory effect on both miRNA-132-3p and TGF-β1 during the committed differentiation of leukemia cells induced by PMA, in order to ascertain whether the miRNA-132-3p/WT1/TGF-β1 signaling axis is involved in the process of differentiation of leukemia cells into macrophages.

## Materials and methods

### Materials

Three human leukemia cell lines (including the acute monocytic leukemia cell lines U937 and THP-1 and the chronic myelocytic leukemia cell line K562; Shanghai Cell Bank, Chinese Academy of Sciences) were employed for the following experiments. PMA (Sigma-Aldrich; cat. no. P1585-1MG) was used as the differentiation inducer. FBS, the BCA protein assay kit and SDS-PAGE gel preparation kit were obtained from Shanghai Biyuntian Biotechnology Co., Ltd. HyClone^®^ RPMI-640 Complete™ medium was purchased from Cytiva. Wright-Giemsa staining solution, the secondary antibody, RIPA protein lysis solution and 5X protein loading buffer were purchased from Beijing Solarbio Science & Technology Co., Ltd. The standard protein marker and the Lipofectamine^®^ 2000 transfection kit were purchased from Thermo Fisher Scientific, Inc. The ECL luminescence kit was purchased from Shandong Sparkjade Scientific Instruments Co., Ltd. Regarding the antibodies, WT1 antibody (cat no. 83535S) was purchased from CST Biological Reagents Co., Ltd., β-actin (cat. no. 20536-1-AP) primary antibody was from Proteintech Group, Inc., TGF-β1 antibody (cat. no. YT4632) was purchased from Immunoway Biotechnology, Inc., and HRP-labeled rabbit secondary antibody (cat. no. ZB-2301) was provided by OriGene Technologies, Inc. Sangon Biotech Co., Ltd. designed and synthesized the primers. The Reverse transcription (RT) kit, PrimeScript™ RT kit with gDNA eraser (for perfect real-time experiments), chimeric fluorescence detection kit and TB Green Premix Ex Taq™ (Tli RNaseH Plus) kit were obtained from Takara Biomedical Technology (Beijng) Co., Ltd. Cell cycle analysis kit was purchased from Jiangsu KGI Biotechnology Co., Ltd. The fluorescent binding antibodies PE-CD11b (cat. no. 301306) and FITC-CD14 (cat. no. 301804) were purchased from BioLegend, Inc. The TRIzol^®^ reagents, which were manufactured by Thermo Fisher Scientific, Inc., were purchased from Tiangen Biotech Co., Ltd. Finally, trichloromethane was purchased from Sinopharm Chemical Reagent Co., Ltd. Cell counting kit-8 (CCK8) was purchased from DOJINDO (Cat.no.CK04). Inhibition of proliferation of leukemia cells induced by PMA. THP-1, U937 and K562 cells were grown in culture flasks containing 10% FBS in RPMI-1640 Complete™ medium, cultured at 37°C in an atmosphere containing 5% CO2. When the three types of cells had reached the exponential growth phase, they were collected and added to the 96-well plates, and the PMA solution was also added, so that the final cell concentration in each well was 1x10^5^/ml, and the corresponding concentration of PMA solution was 100 ng/ml (based on the IC_50_ concentration, where the IC_50_ value represents half-maximal inhibitory concentration). An equal volume of 100 µl RPMI-1640 medium was used as the negative control group. Three duplicate wells were set up for each group, and the three types of cells were subsequently cultured at 37°C in an atmosphere of 5% CO_2_ and with saturated humidity. After 48 h, 10 µl CCK8 solution was added to each well and incubated at 37°C for a further 2 h, and then the OD values of the 96-well plate at 450 nm were measured.

### Wright-Giemsa staining to observe morphology of the THP-1 cells

After the THP-1 cells were induced with 100 ng/ml PMA *in vitro* for 48 h, the morphological changes of the cells were observed directly under an inverted optical microscope, and Wright-Giemsa staining was employed to stain the cells both before and after their exposure to PMA. The main steps of the procedure were as follows: First, 2x10^5^ THP-1 cells of the above-mentioned control and experimental groups were collected after exposing the cells to PMA for 48 h. The cells were subsequently washed twice with cold PBS and resuspended in 100 µl PBS, preparing a cell suspension. The cell suspension was centrifuged at 205 x g for 3 min at 4°C, after which the centrifuged smear was dried and stained with Wright-Giemsa staining solution at room temperature for 5 min. Finally, changes in cell morphology were observed under an optical microscope, and the resulting images were capturedwith full field photography.

### Flow cytometry to detect the expression of the CD11b and CD14 antigens

The THP-1 cell suspension was collected for the control group and different experimental groups, placed into test tubes, and the cells were then resuspended in PBS buffer, before washing twice with PBS and centrifuging the cell suspensions at 205 x g for 3 min at 4°C. The cell concentration was adjusted to 1x10^6^/ml, and 100 µl PBS was added into each test tube. Subsequently, 5 μl PE-labeled mouse anti-human CD11b fluorescent antibody (cat. no. 301306; 1:1,000 dilution; BioLegend, Inc.) or FITC-labeled mouse anti human CD14 fluorescent antibody (cat. no. 301804; 1:1,000; BioLegend, Inc.) was added into each tube, respectively. The cells were incubated in the dark at 4°C for a further 30 min, and then resuspended with 200 μl PBS and fixed in 1% paraformaldehyde at 4°C for 20 min. A FACSVerse flow cytometer (BD Biosciences) was used to measure the expression levels of the CD11b and CD14 antigens. Isotypic rat IgG (BioLegend, Inc.) was used to examine for nonspecific binding. All experiments were repeated three times, and the expression of CD11b and CD14 was evaluated as the mean ratio.

### Western blot assay to detect expression of WT1 and TGFβ1

Chiefly, 3x10^6^ THP-1 cells were first collected from the control group and different experimental groups, an appropriate amount of pre-cooled PBS buffer was added, and the cell mixture was centrifuged at 462 × g for 5 min at room temperature, before washing the cells twice with PBS buffer. After discarding the supernatant, the cells were lysed using RIPA buffer (Beijing Sun Biotechnology Co., Ltd.) to extract total cell proteins, and the protein concentration was then determined using the BCA method. Subsequently, 5X protein loading buffer was added, and the proteins were denatured by boiling at 95°C for 10 min. Aliquots (20 μg) of protein were then loaded on to SDS-PAGE gels to perform SDS-PAGE (10% gels), and the separated proteins were then transferred onto a PVDF membrane. Subsequently, the membrane was sealed with 5% skimmed milk at room temperature for 90 min, prior to incubation with the corresponding primary antibody. WT1 (1:1,000), TGFβ1 (1:10,000) and GAPDH (1:2,000) antibodies were added, and the primary antibodies were incubated with the membrane overnight at 4°C. The membranes were washed with 1X TBST for 30 min. Subsequently, after the first antibody reactions were completed, the membranes were exposed to goat anti-rabbit IgG secondary antibody (1:20,0000 dilution) for an incubation at room temperature for 90 min. Finally, the membranes were washed with 1X TBST for 30 min. The proteins were detected using an ECL kit (Sparkjade ECL Super; cat no. ED0015-B; Shandong Sparkjade Scientific Instruments Co, Ltd,), and ImageJ, 1.51j8 software was used to determine the densities of the protein bands. All experiments were repeated three times, and the results were expressed as the average relative expression densities.

### Databases used to predict the association between miRNA-132-3p and WT1

Three databases (TargetScan (http://www.targetscan.org), PITA(http://genie.weizmann.ac.il/ pubs/mir07/mir07_data.html) and microRNAorg (http://www.microrna.org/), were selected to accurately predict the possible upstream miRNAs of WT1. Based on the potential miRNAs predicted by each database, a cross-analysis of the three databases was conducted. Through an analysis of the intersection of the three databases, common target miRNAs were obtained, and the most likely potential miRNAs to target WT1 were selected on the basis of the size of the P-value. In addition, differential miRNAs between the PMA experimental group and negative control group were detected and analyzed using the GeneChip miRNA 4.0 detection chip (Affimatrix Co., Ltd.) The types of miRNAs that were upregulated after induction of differentiation were screened based on their P-values. In addition, the abovementioned databases were also used to predict miRNAs that were associated with macrophage differentiation. The upregulated miRNAs identified by crosstalk from the mentioned three different analyses were selected to represent the potential miRNAs for targeting WT1.

### RT-quantitative (RT-q) PCR assay to detect the expression of miRNA-132-3p and WT1

The THP-1 cells were observed under an inverted microscope until they had reached the exponential phase of growth. Subsequently, 1x10^6^ cells were collected, and the total RNA was extracted using TRIzol^®^, following the instructions provided by the kit (Thermo Fisher Scientific, Inc.). After the total RNA concentration had been measured using a Ultra-micro Nucleic Acid Protein Analyzer, the cDNA was synthesized using an RT kit [specifically, according to the instructions of the PrimeScript^®^ RT kit and the gDNA Eraser (Perfect Real Time) RT kit]. The main steps of the RT reaction were carried out; for example, the sample was incubated in the Eppendorf PCR system at 42°C for 15 min, after which it was incubated at 85°C for 5 secmin, and then incubated at 4°C for a further 2 min. The cDNA obtained by RT was then used as the next PCR amplification template. The sequence of the miRNA-132-3p sense strand was 5’-CGATACCGTTCTAACAG TCTACAGC-3’, whereas that of the antisense strand was 5’-TATGGTTTTC ACGACTGTGTGAT-3’; the sequence of the U6 sense strand was 5’-CTCGCTTC GGCAGCACA-3’, and that of the antisense strand was 5’-AACGCTTCACGAATT TGCGT-3’, and these primers were synthesized by Shanghai GenePharma Co, Ltd. These two primers were designed according to the stem-loop method. The downstream and RT primers of the stem-loop method were artificially added (Shanghai Sangon Biotech, Co., Ltd.). The WT1 sense strand was 5’-CAGGCTGCAATAAGAGATATTTTAAGCT-3’, and the antisense strand was 5’-GAAGTCACACTGGTATGGTTTCTCA-3’; the GAPDH sense strand was 5’-CAACTTTGGTATCGTGGAAGG-3’, and the antisense strand was 5’-GCCATCACGCCACAGTTTC-3’, and these were synthesized by Biosune Biotechnology (Shanghai) Co., Ltd. The real-time fluorescence quantitative amplification reactions were performed mainly following the instructions of the TB Green^®^ Premium Ex Taq (Tli RNaseH Plus) kit. The thermocyling conditions of the RT-qPCR process mainly comprised the following key steps: 10 sec at 95°C; 5 sec at 60°C, and 10 sec at 72°C for a total of 40 cycles, with a final step of 60°C for 34 sec. The results were quantified with the 2^-Δ Δ Cq^. These experiments were repeated three times, and the results were expressed as the mean ± standard deviation.

### Dual-luciferase reporter gene assay for miRNA-132-3p targeting of WT1

The regional sequence of WT1 gene for miRNA-132-3p binding was predicted using the ‘TargetScan’ bioinformatics website (http://www.targetscan.org). It was entrusted to Biosune Biotechnology (Shanghai) Co., Ltd. to construct the wild-type plasmid vector pmirGLO-WT1-wt, and the mutant plasmid vector pmirGLO-WT1-mut (www.Biosune.com). 293T cells (as a density of 7x10^4^ cells/well) were added to the wells of a 24-well culture plate, and subsequently incubated for further 24 h at 37°C in an atmosphere of 5% CO_2_ and with saturated humidity. Following the instructions provided in the cell transfection kit (jetPRIME; Polyplus Co., Ltd), the WT1 wild-type plasmid, the WT1 mutant plasmid and miRNA-132-3p mimics were co-transfected into the 293T cells. After further cultivation for 48 h, the luciferase activity was measured using a dual-luciferase reporter assay system (EnVision™; PerkinElmer Co., Ltd.), according to the manufacturer’s instructions. Data were normalized for transfection efficiency by dividing the firefly luciferase activity with that of *Renilla* luciferase.

### Cell transfection assay

Cell transfection reactions were performed following the instructions provided by the kit. For the short interfering RNA (siRNA) interference experiment, FAM fluorescence-labeled si-NC was used as the negative control in parallel to determine the transfection efficiency. According to the instructions of the transfection kit, the siRNA was mixed with the transfection reagent to form a transfection complex. At the same time, the corresponding negative control siRNA NC was also transfected. After that, their transfection complexes were added the inoculated cells in a 6-well plate, and the cells were then cultured for 48 h at 37°C for the subsequent associated detection experiments. The sequence of the si-WT1 sense strand was 5’-CUACAGCAGUGA CAAUUUATT-3’, whereas that of the si-WT1 antisense strand was 5’-UAAAUUGU CACUGCUGUAGTT-3’. The sequence of the si-NC sense strand was 5′-UUCUCCGA ACGUGUCACGUTT-3′, and that of its antisense strand was 5′-ACGUGACACGUU CGGAGAATT-3′. The siRNAs were synthesized by Shanghai GenePharma Co., Ltd.

Regarding the siRNA interference experiments, four experimental groups were set up to perform the siRNA interference experiments, as follows: the 0 μg/l control group, the 100 μg/l PMA experimental group, the 100 μg/l PMA+si-NC transfection experimental group, and the 100 μg/l PMA+si-WT1 transfection experimental group. The si-WT1 transfer methods were followed as per the instructions provided with the transfection kit. In addition, the role of the miRNA-132-3p mimic in modulating WT1 expression was also investigated. The sequences of the primers for the miRNA-132-3p mimic were as follows: Sense, 5’-UAACAGUCUACAGCCAUGGUCG-3’; and antisense, 5’-CGACCAUGGCUGUAGACUGUUA-3’; and the sequence of the mimic NC sense strand was 5’-UUCUCCGAACGUGUCACGUTT-3’, whereas that of the antisense strand was 5’-ACGUGACACGUUCGGAGAATT’3’. These primers were also synthesized by Shanghai GenePharma Co, Ltd., and the transfection experiments were performed using Lipofectamine^®^ 2000, following the instructions provided by the kit. miRNA-132-3p mimic was transfected into the THP-1 cells, and RT-qPCR and western blotting were performed to detect the expression levels of miRNA-132-3p and WT1 protein, respectively, as described above. At the same time, the corresponding expression levels of CD11b and CD14 of THP-1 cells were detected by flow cytometric assay.

### Chromatin immunoprecipitation (ChIP) assay

ChIP assay was performed to detect the targeted binding and regulation of TGF-β1 by WT1. After induction by PMA for 2 h, a total of 2x10^7^ THP-1 cells were collected, and subsequently formaldehyde at a final concentration of 1% was added into the tubes, and the mixture was incubated further for approximately 10 min. After the preparation of cross-linked chromatin had been completed, ultrasound treatment was used to achieve an average size of DNA fragments of 200-1,000 bp, as determined by ethidium bromide gel electrophoresis. The purified chromatin was immunoprecipitated using 3 μg WT1-specific antibodies or an irrelevant antibody (IgG). The immunoprecipitates were kept overnight at a temperature of 4℃. Finally, the immunoprecipitated chromatin was used as the template for amplification by PCR. Annealing reactions were performed at 59.2°C, and the PCR products are separated by routine agarose gel electrophoresis. The primers for TGF-β1 were as follows: Sense, 5’-ATTAAGCCTTCTCCGCCTGGTCCT-3’, and antisense, 5’-CGGCAACGGAAAA GTCTCAAAAGT-3’ [Biosune Biotechnology (Shanghai) Co, Ltd.].

### Rescue experiment of miRNA-132-3p by si-TGF-β1 upon inducing differentiation of the THP-1 cells

Regarding the si-TGF-β1 interference experiments for reversing the effects of miRNA-132-3p in inducing differentiation of the THP-1 cells, three experimental groups were set up; namely, the miRNA-132-3p mimics-NC, miRNA-132-3p mimics and miRNA-132-3p mimics + siTGF-β1 groups. Following the instructions provided by the manufacturer of the kit for the Lipofectamine 2000™ transfection assay, the miRNA-132-3p mimics, siTGF-β1 and their si-NCs were transfected into the THP-1 cells. At 20 h post-transfection, the cells were collected, and the expression of TGF-β1 protein was detected by western blotting assay, whereas the expression levels of CD11b and CD14 were detected by FACS at the same time. We have designed four pairs of si-TGFβ1, and the interference sequence was validated for interference efficiency through qRT-PCR and Western blotting experiments, and the pair of sequences with the most obvious interference efficiency was selected as the candidate siRNA in the present manuscript.The sequence of this candidate siTGF-β1 sense strand was 5′-CACUGCAAGUGGACAUCAATT-3′, whereas that of the antisense strand was 5’-UUGAUGUCCACUUGCAGUGTT-3′. The si-NC sense strand was 5′-UUCUCCGAACGUGUCACGUTT-3’, whereas that of the antisense strand was 5’-ACGUGACACGUUCGGAGAATT-3’ (Shanghai GenePharma Co., Ltd.). The sequences of the miRNA-132-3p mimics-NC and miRNA-132-3p mimics primers were the same as mentioned above.

## Statistical analysis

The data were processed using GraphPad Prism 8 software, and the quantitative data were analyzed using Shapiro Wilk (S-W) normal distribution. The differences between two groups were compared using independent sample t-tests. ANOVA analysis was used for making comparisions with multiple groups. One-way ANOVA was performed using the Bonferroni test. Pearson correlation analysis was performed to assess the correlation between miR-132-3p and WT1. Double-tailed tests were also employed for analysis of all the data. Each experiment was repeated three times, and the results were expressed as the mean ± standard deviation. P<0.05 was considered to indicate a statistically significant difference.

## Results

### Cell growth density and proliferation of leukemia cells induced by PMA

Induced by PMA at a concentration of 100 ng/ml, the K562, U937 and THP-1 cells were all found to be significantly inhibited by PMA as compared with that of the respective control groups (0.67±0.01 vs. 0.43±0.02, t=22.88, P<0.001; 0.67±0.01 vs. 0.34±0.02, t=28.81, P<0.0001; and 0.67±0.01 vs. 0.32±0.02, t=17.22, P<0.0001) (Fig. 1A). Based on the fact that the most significant inhibitory effects on cell proliferation were observed with the THP-1 cell line, THP-1 cells were selected to construct a model of committed differentiation from leukemia cells into macrophages, and these were subsequently used for the further experiments. As shown in Fig. 1B and C, after exposure to PMA for 48 h, the cell density of the THP-1 cells was significantly reduced as observed under a microscope, and the proliferation of the THP-1 cells was significantly inhibited by PMA (0.65±0.03 vs. 0.47±0.01, t=8.522, P<0.01). The results of the flow cytometry analysis showed that the G0/G1 phase of THP-1 cells increased from 41.07 to 47.38% (41.04±1.05% vs. 47.38±0.42%, t=9.678, *P*=0.0006), whereas that of the S phase decreased from 41.85 to 26.27% (41.85±3.49% vs. 26.27±1.16%, t=7.337, *P=*0.0018), and the G2/M phase was increased from 17.08 to 26.35% (17.08±2.44% vs. 16.735±1.19%, t=5.908, *P*=0.0041). Taken together, these experiments showed that the proliferation of THP-1 cells could be inhibited by PMA, and that PMA could induce cell cycle arrest at G0/G1 phase and G2/M phase (Fig. 1D-F).

**Fig. 1.**
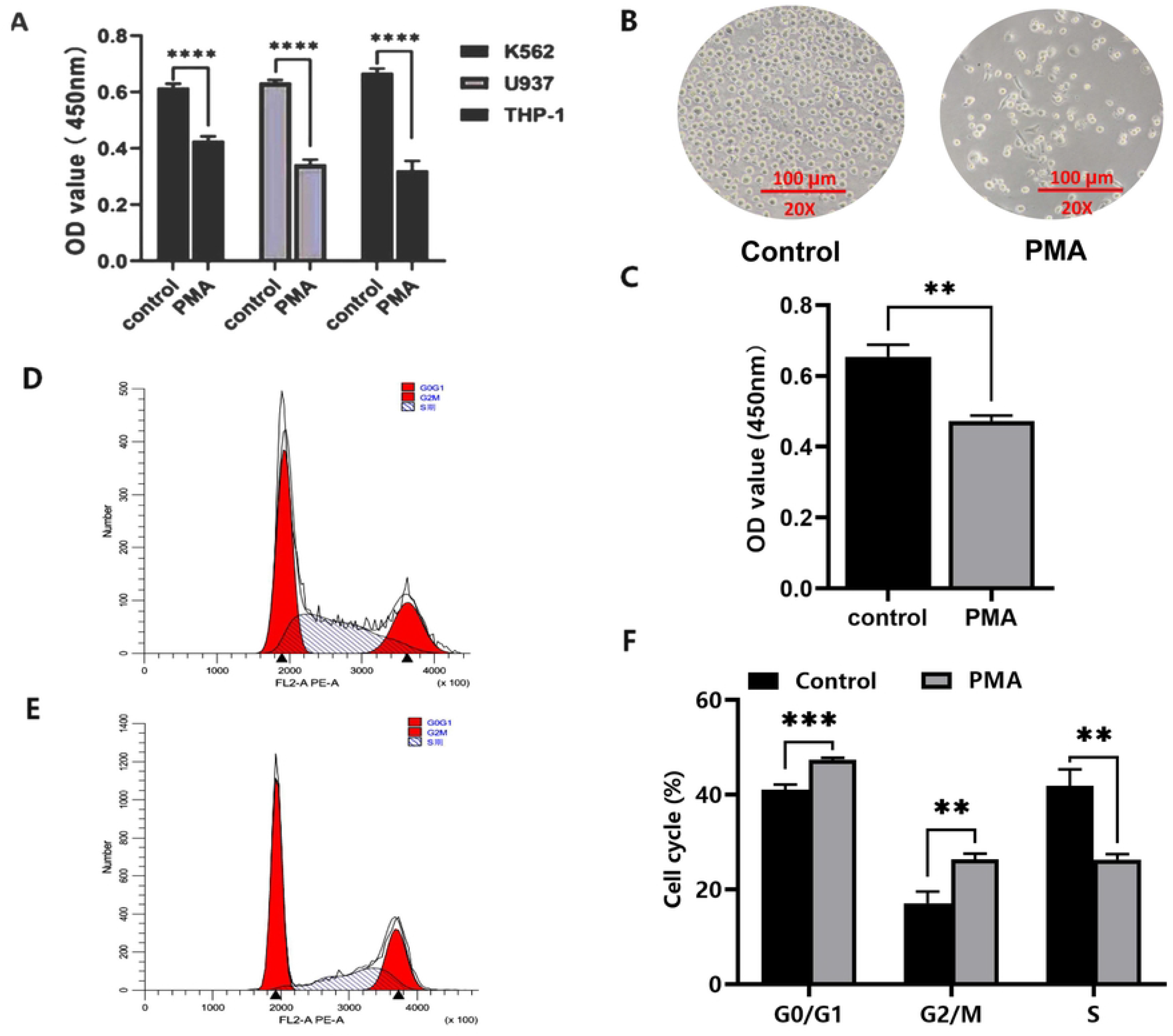
The proliferation and cell cycle analysis of leukemia cells after exposure to PMA K562, U937 and THP-1 leukemia cells were treated with PMA 100 ng/mL for 48 hours, the cell density were directly observed under a microscope, and proliferation of these leukemia cells were detected by CCK8 cell counting assay. Cell cycle was detected by Flow cytometry assay. A: Proliferation of K562, U937 and THP-1 leukemic cells before and after exposure to PMA; B. Cell density of THP-1 cells under microscopy; C: CCK8 assay to detect the proliferation of leukemia THP-1 cells; D. Cell cycle changes of control group THP-1 cells by flow cytometry assay; E: Cell cycle changes of PMA group cells by flow cytometry assay; F: The average percentage of cell cycle distribution. (* *P* < 0.05, ** *P* < 0.01, **** *P* < 0.0001)

### Expression of the CD11b and CD14 differentiation antigens of the leukemia THP-1 cells

On the basis of the cell proliferation experiments which confirmed that PMA could significantly inhibit the proliferation of THP-1 cells, further experiments were performed to detect the committed differentiation level of this type of cells into macrophages. Usually, in the committed differentiation experiment of leukemia cells into monocytes and macrophages, the quantitative detection of classic CD11b and CD14 differentiation antigens is mainly used. However, in order to more intuitively demonstrate the successful model of leukemia cells to differentiate into monocytes and macrophages, we also conducted Wright-Giemsa staining in the differentiation experiment induced by PMA. As compared with the control group (Fig. 2A), after exposure to PMA *in vitro* for 48 h, the THP-1 cells were observed to have increased in size, and the ratio of cytoplasm to nucleus was increased, moreover, certain cells exhibited two or even multiple nuclei, showing a trend of differentiation towards macrophages (Fig. 2B), and the population of more matured differentiated THP-1 cells had clearly increased (Fig. 2C) (*P*<0.001). The expression levels of the CD11b and CD14 antigens, which acted as macrophage-specific markers on the cell surface of the THP-1 cells, were also detected by flow cytometry. The expression of CD11b was found to be upregulated to a greater extent compared with that in the control group (36.09±2.63% vs. 2.63±0.05%, t=30.34, *P*<0.0001) (Fig. 2D and E), and the expression of CD14 was also higher compared with that in the control group (26.93±2.34% vs. 3.60±0.52%, t=68.73, *P*<0.0001) (Fig. 2F and G). Therefore, both examining the morphological changes into more mature cells, and detecting the upregulation of CD11b and CD14 expression served to demonstrate that THP-1 cells could be induced by PMA towards differentation into macrophages.

**Fig. 2.**
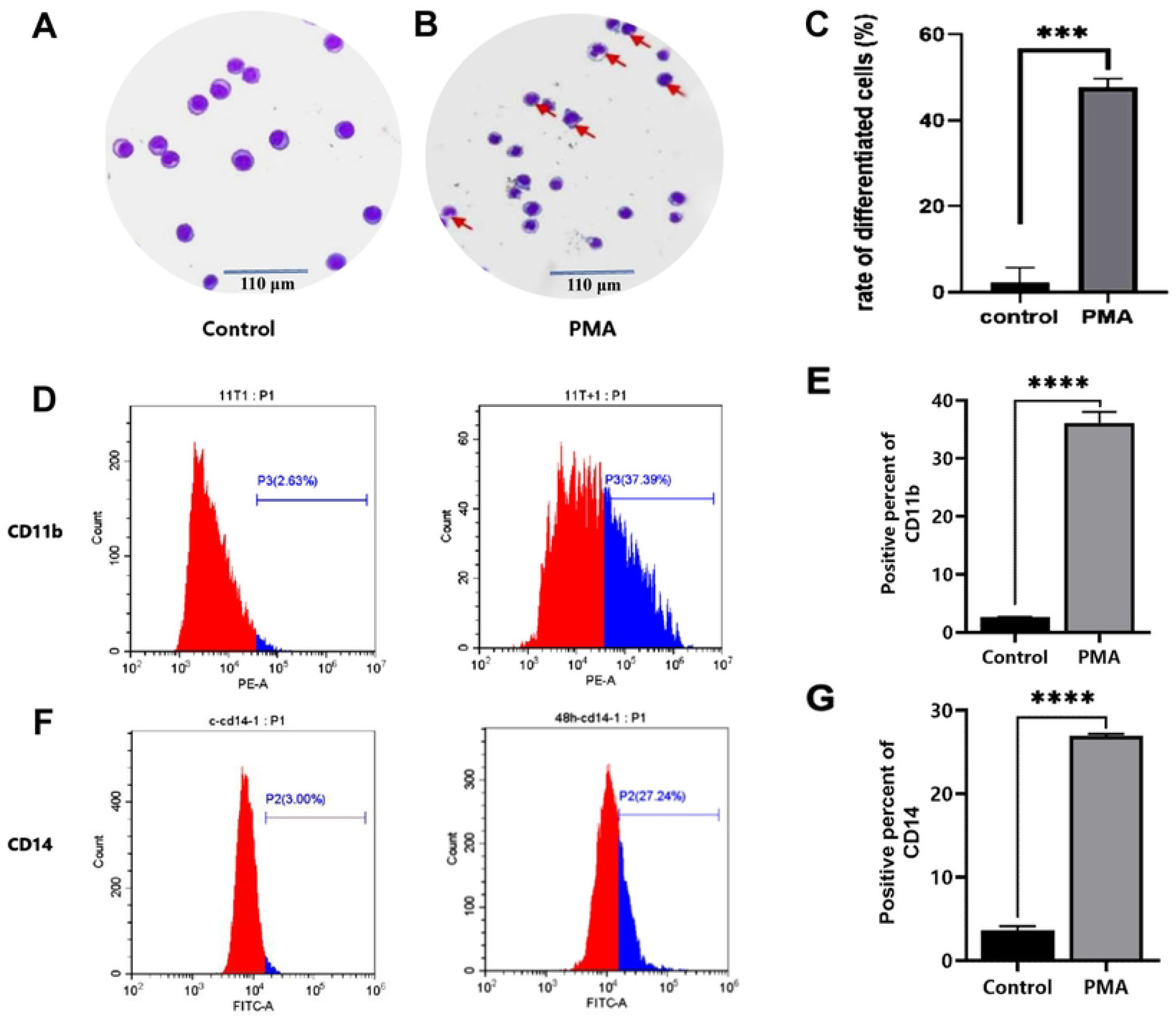
Effects in differentiation of THP-1 leukemia cells after PMA treatment Induced by PMA for 48 hours, the THP-1 cells were collected and stained with Wright-Giemsa staining, and expression of CD11b and CD14 antigens on the cell surface was detected by flow cytometry. A. Wright-Giemsa staining of THP-1 cells control group; B. Wright-Giemsa staining of THP-1 cells induced by PMA; C. The mean rate of differentiated THP-1 cells; D: Expression changes of CD11b expresson percent in THP-1 cells in control group and PMA-induced group by flow cytometry assay; E. Mean expression of CD11b in control group and PMA-induced group THP-1 cells; F. Expression changes of CD14 expresson percent in control group and PMA-induced group by flow cytometry assay; G: Mean CD14 expression percent in control and PMA-induced group. (**** *P* < 0.0001)

### Role of WT1 in PMA-induced committed differentiation of THP-1 cells

In our pre-experimental studies, the KEGG and BIOCARTA databases were used to analyze and screen genes associated with PMA-induced cells, and a Pubmed/MEDLINE literature research was performed to identify the oncogenes that were associated with proliferation of the leukemia cells. After crosstalk analysis, it was found that WT1 was involved in the proliferation of leukemia cells, and therefore our attention turned to detecting the role of WTI in regulating the differentiation of THP-1 cells into macrophages. These experiments revealed that the expression of WT1 protein in the THP-1 cells was downregulated following exposure to PMA at the indicated time point (1.05±0.06 vs. 0.54±0.05, t=11.18, *P*=0.0004) (Fig. 3A and B). Subsequently, the potential role of WT1 vector in reversing the effects mediated by the inducer PMA was investigated, and it was found that the level of WT1 was significantly reduced following PMA induction; however, when transfected with WT1 vector at the same time, the effect of PMA on reducing WT1 expression was significantly reversed by WT1 vector. Compared with the PMA group, the expression level of WT1 in the group transfected with both PMA and the WT1 vector was significantly increased (0.55±0.09 vs 0.96±0, 13, t=4.38, *P*=0.012) (Fig. 3C and D).

**Fig. 3.**
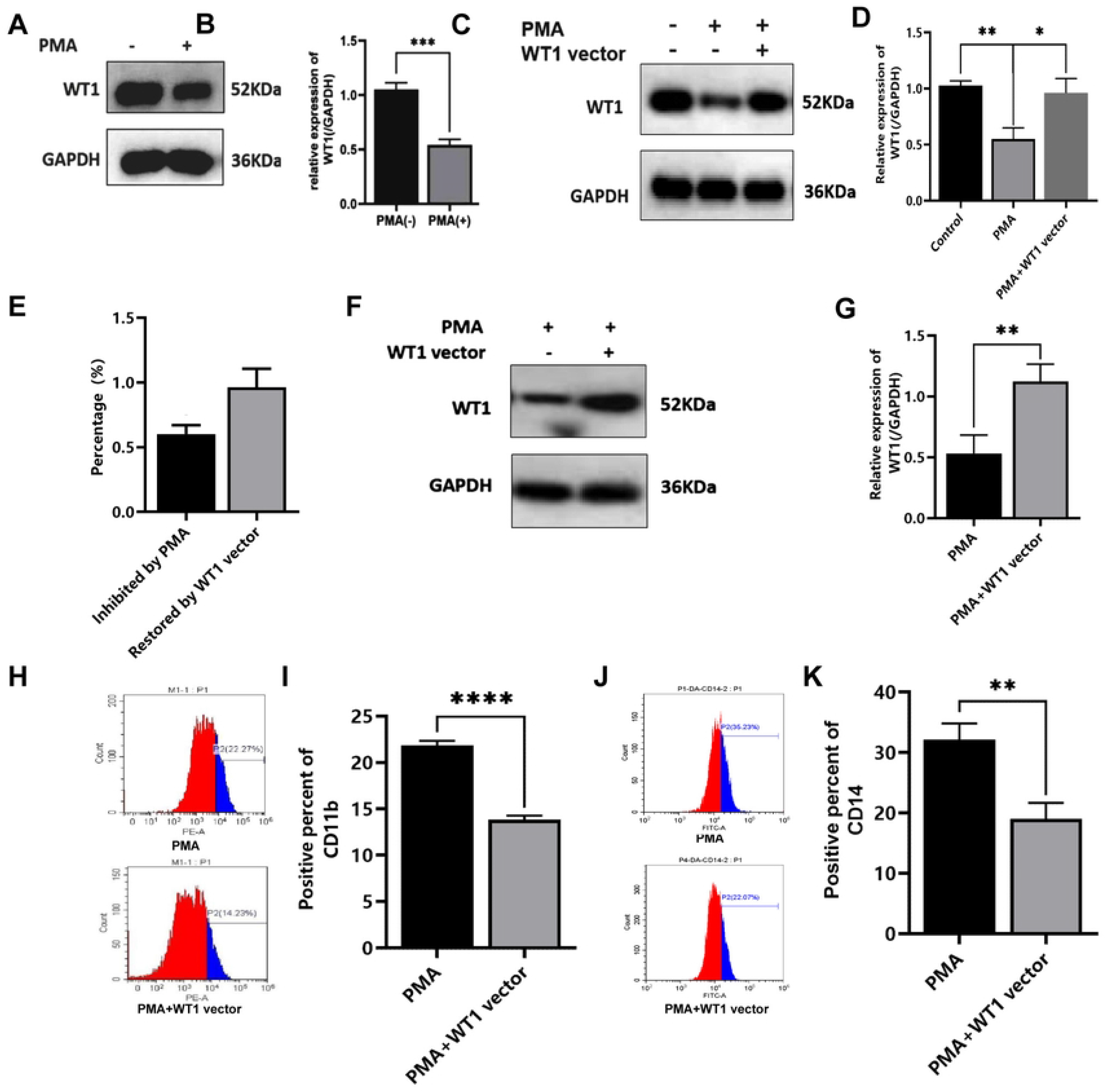
Effect of WT1 gene overexpression on THP-1 cell differentiation induced by PMA Induced by PMA for 48 hours, the THP-1 cells were collected, the expression of WT1 cells was detected by western blotting assay, and the CD11b and CD14 of THP-1 cells was detected by flow cytometry. A. WT1 protein expression in THP-1 cells induced by PMA; B: Mean WT1 protein expression of THP-1 cells induced by PMA. C: WT1 expression by western blotting in THP-1 cells transfected with WT1 vector as compared with control group or PMA group; D: Mean relative expression of WT1 in THP-1 cells transfected with WT1 vector as compared with control group or PMA group; E. The inhibition percent of WT1 expression by PMA and the restoration percent of WT1 expression by PMA+WT1 vector to the control group, which were based on the results of Figure 3C and D. F. The WT1 expression by Western blot assay; G. Mean relative expression of WT1 in THP-1 cells transfected with WT1 vector. H. The CD11b expression by flow cytometry assay; I. Effect of WT1 overexpression by transfection with WT1 vector on CD11b differentiation antigen induced by PMA; J. The CD14 expression by flow cytometry assay; K. Effect of WT1 overexpression by transfection with WT1 vector on CD14 differentiation antigens induced by PMA. G: WT1 protein expression of THP-1 cells transfected with WT1 vector. ;H: (* *P <* 0.05, ** *P <* 0.01, *** *P <* 0.001)

When compared with the control group, the mean rate of decrease in the expression level of WT1 mediated by PMA was 60.20±6.93%, whereas the mean rate of increase in WT1 expression for the WT1 vector + PMA group was 96.29±14.36% (Fig. 3E). To further detect the role of WT1 overexpression in inducing THP-1 cells towards differentiation into macrophages, THP-1 cells were treated with WT1 vector when exposed to PMA. It was found that the WT1 vector reversed the decrease in WT1 expression that was observed upon induction by PMA (0.53±0.15 vs. 1.12±0.14, t=4.928, *P*=0.0079) (Fig. 3F and G). The effect of WT1 overexpression on the differentiation of THP-1 cells induced by PMA was subsequently assessed by investigating the levels of the CD11b and CD14 antigens by flow cytometric analysis, and this revealed that the percentages of either CD11b- or CD14-positive cells in the WT1 vector + PMA group were downregulated compared with those in the PMA-induced group alone (13.85±0.43% vs. 21.83±0.52%, t=20.61, P<0.0001; and 19.06±2.61% vs. 32.13±2.69%, t=6.028, P=0.0038, respectively) (Fig. 3H and I). Taken together, these results suggested that overexpression of WT1 is able to counteract the differentiation-induced effects on the THP-1 cells mediated by PMA.

### Cross analysis of potential miRNAs targeting WT1, and their associations during committed differentiation towards macrophages

The putative targets of miRNA-132-3p were predicted by analysis of the TargetScan (http://www.targetscan.org/) and miRTarBase (http://mirtarbase.mbc.nctu.edu.tw/php/index.php) databases, also with the aim of predicting potential upstream miRNAs that were associated with WT1. The prediction results indicated that there were 17 miRNAs that could target and regulate WT1 (Fig. 4A). To investigate whether differential changes occurred to the miRNAs in the THP-1 cells upon induction by PMA, RNA sequencing (RNA-seq) technology was used to identify differentially expressed genes and upregulated miRNAs, and a difference multiple >2 and a P-value <0.05 were taken as the boundaries. Through visual clustering analysis, it was found that there were significant increases or decreases in miRNAs both before and after PMA induction, and an MA plot also revealed the same increases or decreases of the miRNAs. A total of 69 miRNAs that were differentially expressed in THP-1 cells induced by PMA were identified (Fig. 4B). The MirDB and TargetScan databases were also utilized to predict the miRNAs associated with macrophage differentiation, and upon analyzing the intersection of the screening results of the two databases, the results indicated that 56 miRNAs were associated with monocytes/macrophages (Fig. 4C). Subsequently, the abovementioned miRNAs that could target WT1, the miRNAs associated with macrophage differentiation according to the database analysis, and the differentially expressed miRNAs before and after PMA induction were analyzed by intersection analysis, and these results indicated that only miRNA-132-3p and miR-212-3p were highly correlated with WT1 expression (Fig. 4D). Further experiments revealed that, as compared with the control group, the expression of miRNA-132-3p in the THP-1 cells was significantly upregulated (3.54±0.09 vs. 0.36±0.03, t=55.02, P<0.0001) (Fig. 4E). However, by contrast, the expression of miR-212-3p remained almost unchanged before and after the induction of differentiation (0.43±0.02 vs. 0.43±0.04, t=0.3462, P>0.05) (Fig. 4F). Collectively, these results suggested that miRNA-132-3p may be an upstream miRNA associated with targeting the regulation of WT1, although further validation experiments need to be performed in this regard.

**Fig.4.**
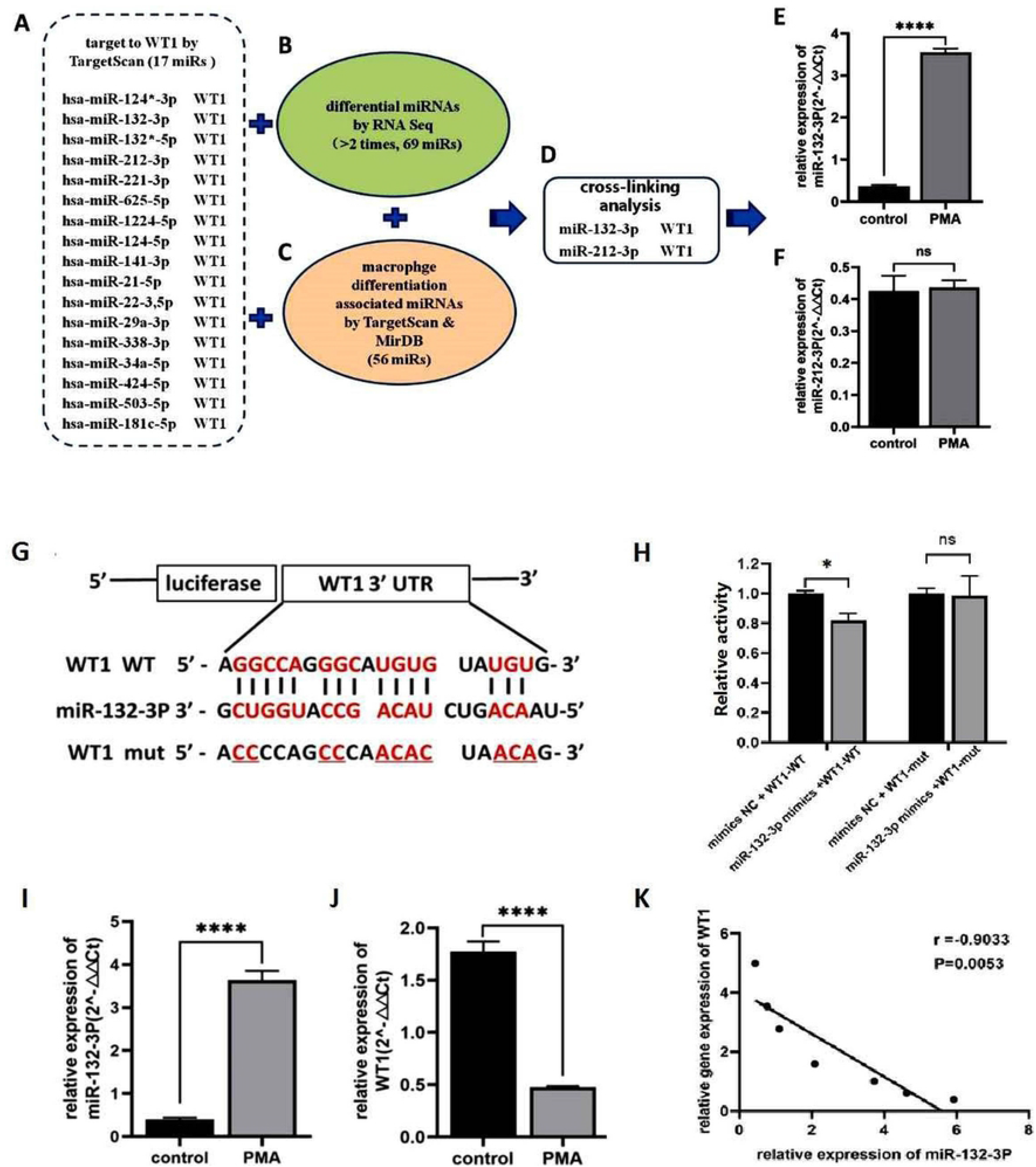
Potential miRNA targeting WT1 and their relationship during committed differentiation towards macrophages TargetScan and miRTarBase were used to predict potential upstream associated miRNAs of WT1. RNA seq technology to identify differentially altered genes (greater than 2 and a *P*<0.05) as the boundary. TargetScan and MirDB databases were used to predict the miRNAs associated with monocyte/macrophage differentiation, and the intersection of the two databases was tried. Further validation of positively related miRNAs was demonstrated by qRT-PCR assay. In order to further verify whether miRNA-132-3p could bind WT1 3’UTR and modulate its promoter activity, a dual luciferase reporter assay was used. The relative expression of miRNAs was detected by qRT-PCR assay, and the correlation analysis between expression of miRNA-132-3p and WT1was indicated by Pearson analysis. A. TargetScan database predicts the possible miRNAs that target and bind to WT1; B. The number of differential miRNA species before and after differentiation and the number of macrophage differentiation-associated miRNA species analyzed by database; C. Target gene prediction, differential detection before and after differentiation, and database analysis macrophage-associated miRNA intersection analysis; D. miRNA-132-3p and miR-212-3p related to WT1 expression by intersection analysis; E: related expression of miRNA-132-3p before and after differentiation. F: miR-212-3p expression before and after differentiation; G: miRNA-132-3p targets WT1 3̛ UTR sequence prediction; H. Luciferase activity assay of miRNA-132-3p targeting WT1; I. The relative expression level of miRNA-132-3p in THP-1 cells before and after PMA induction; J. Relative expression levels of WT1 in THP-1 cells before and after PMA induction; K. Correlation analysis between miRNA-132-3p and WT1 expression. (ns: no significant*, * P<0.05, \**\*\*\* *P* < 0.0001)

Furthermore, dual-luciferase reporter assay experiments were performed to detect the targeted binding of miRNA-132-3p to the 3’-UTR of WT1. The TargetScan database predicted that miRNA-132-3p may target binding to the 3’-UTR of WT1, although further validation of this point is required. In order to further verify whether miRNA-132-3p is targeted to the WT1 3’-UTR, a dual-luciferase reporter gene experiment was designed. First, the recombinant plasmids pmirGLO hWT1-3’-UTR-MUT and pmirGLO hWT1-3’-UTR-WT were constructed, and the WT1 wild-type and WT1 mutant plasmids constructed above were transfected into THP-1 cells, together with miRNA-132-3p mimics or NC mimics, respectively (Fig. 4G). When co-transfected with miRNA-132-3p mimics, the luciferase activity of the pmirGLO hWT1-3’-UTR-WT group was lower compared with that of the co-transfection group of pmirGLO hWT1-3’-UTR-WT including NC mimics (0.82±0.04 vs. 1.00±0.01; t=8.086, P<0.05, Fig. 4H). This difference was found to be statistically significant. However, the luciferase activity of the WT1 mutant plasmid co-transfected with miRNA-132-3p mimics did not change significantly compared with NC mimics (1.00±0.01 vs. 0.98±0.13, t=0.172, P>0.05) (Fig. 4H). These results confirmed that miRNA-132-3p mimics could regulate the expression of WT1 by ‘sponging’ the 3’-UTR of WT1. In addition, the expression of miRNA-132-3p was found to be higher compared with that of the control group following exposure to PMA, and this difference was statistically significant (3.63±0.21 vs. 0.40±0.03, t=25.77, *P*<0.0001) (Fig. 4I). However, the expression level of WT1 was clearly downregulated compared with the control group (0.47±0.01 vs. 1.77±0.09, t=23.13, *P*<0.0001) (Fig. 4J). During the process of differentiation, miRNA-132-3p showed an increase in its expression level, whereas WT1 showed a decrease in expression level. miRNA-132-3p expression was found to be negatively correlated with WT1 expression, and this correlation was statistically significant (Fig. 4K).

### Rescue experiment of miRNA-132-3p targeting WT1 to regulate THP-1 cell differentiation

The above-mentioned results have been shown that downregulation of WT1 expression by PMA can induce differentiation of THP-1 cells, whereas miRNA-132-3p has a negative correlation with WT1. Therefore, a rescue experiment targeting miRNA-132-3p was designed to observe variations in the differentiation of THP-1 cells via targeting WT1. The transfection efficiency of miRNA-132-3p mimics was firstly demonstrated by qRT-PCR (Figure 5A), and the transfection efficiency of WT1 vector was demonstrated by Western blotting experiments and the relative expression of WT1 was indicated as supplimentary Figure 1(supplimentary Figure 1E, F). The results obtained showed that, compared with the mimics NC group, the expression level of WT1 in the miRNA-132-3p mimics-transfected group was clearly downregulated (0.901±0.067 vs. 0.575±0.09, t=5.159, *P*=0.0067) (Fig. 5B and C). Moreover, the expression level of WT1 in the miRNA-132-3p mimics + WT1 vector group was found to be higher compared with that in the miRNA-132-3p mimics group, and the differences were statistically significant (0.58±0.09 vs. 1.04±0.06, t=7.395, *P*=0.0018) (Fig. 5B and C). Subsequently, the levels of the CD11b and CD14 antigens of the THP-1 cells were detected. These experiments showed that the expression of CD11b in the miRNA-132-3p mimics group was higher compared with that in the mimics NC group (22.65±0.27% vs. 3.78±0.32%, t=78.13, *P*<0.001), and the expression of CD14 was also higher in the miRNA-132-3p mimics group (25.01±2.52% vs. 3.55±0.64%, t=14.28, *P*=0.0001). By contrast, the level of CD11b in the miRNA-132-3p mimics + WT1 vector group was lower compared with that in the miRNA-132-3p mimics group (12.77±2.07% vs. 22.65±0.27%, t=8.212, *P=*0.0012) (Fig. 5D, E). The CD14 antigen was also downregulated in the miRNA-132-3p mimics + WT1 vector group compared with that of the miRNA-132-3p mimics group (17.15±0.76% vs. 25.01±2.52%, t=5.170, *P=*0.0067) (Fig.5F, G), and these differences were statistically significant. Taken together, these results suggested that overexpression of miRNA-132-3p led to a downregulation of the expression of WT1 through a negative regulation, thereby promoting the differentation of THP-1 cells towards macrophages.

**Fig.5.**
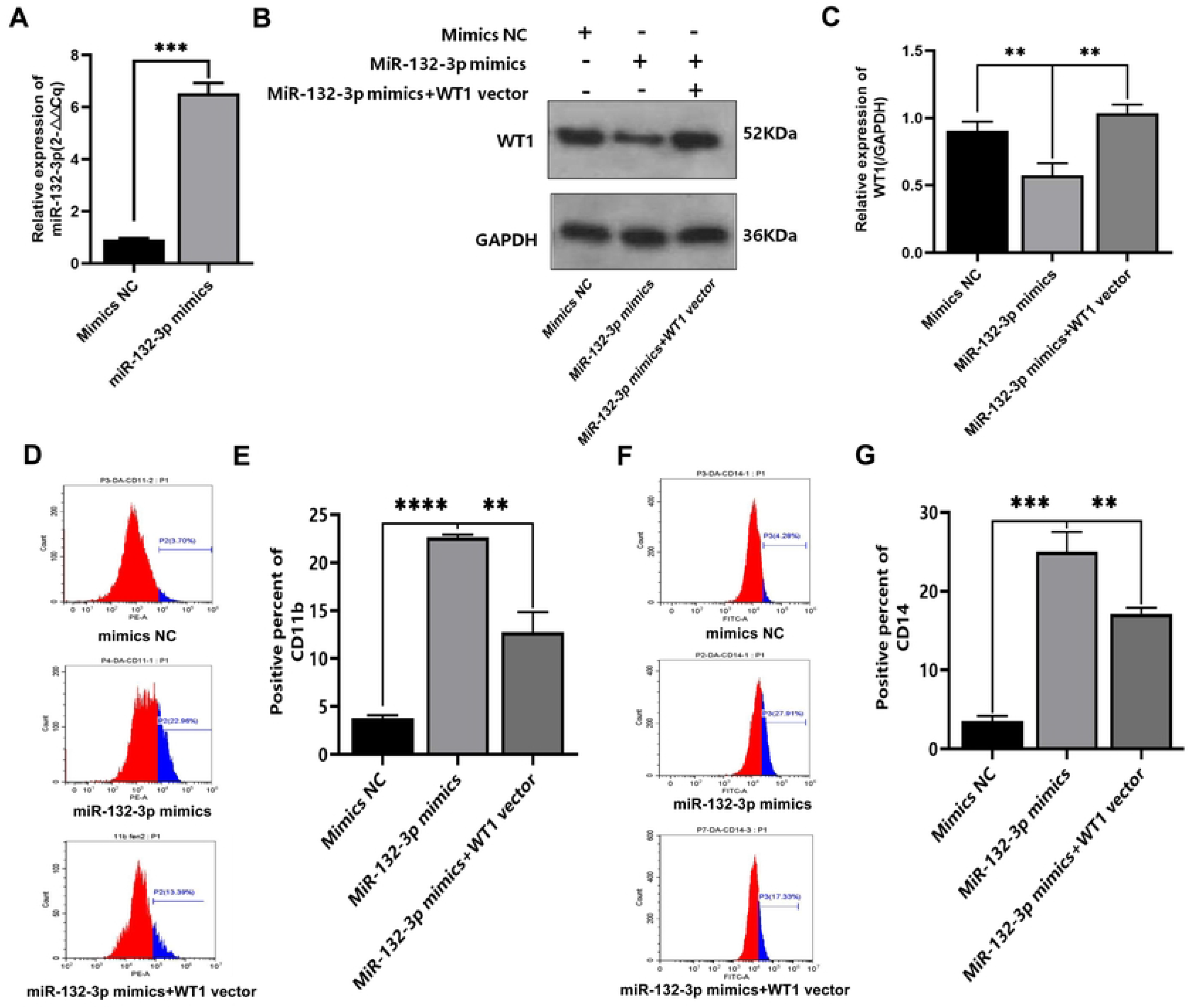
Results of rescue experiments of miRNA-132-3p targeting WT1 to regulate the differentiation of THP-1 cells Rescue experiment about miRNA-132-3p to regulate differentiation of THP-1 cells by targeting WT1 was tried. WT1 protein expression was detected by Western blotting assay, the expression level of CD11b and CD14 differentiation antigen were detected by flow cytometry assay. A. The transfection efficiency of miRNA-132-3p mimics was detected by qRT-PCR; B. WT1 protein expression was detected by Western blot; C. Mean WT1 expression level; D. CD11b expression by flow cytometry; E. Mean expression level of CD11b differentiation antigen; F.CD14 expression by flow cytometry; G. Mean expression levels of CD14 differentiation antigens. (** *P* < 0.01, *** *P* < 0.001)

### Upregulation of TGF-β1 expression by WT1 was detected during differentiation of THP-1 cells towards macrophages

The web tool GeneMANIA (http://genemania.org/) and Alggen’s online database were used to search or predict the association between WT1 and TGF-β1, and to investigate whether WT1 as a transcription factor can target the regulation of TGF-β1. Searching of the Alggen online database revealed that the TGF-β1 promoter region from -180 to +1 bp contains possible binding sites for WT1, suggesting that WT1 may be closely associated with TGF-β1. Upon induction by PMA, western blot analysis was first used to detect TGF-β1 expression of the THP-1 cells, and the results obtained showed that there was an increase in TGF-β1 expression following exposure to PMA (0.379±0.063 vs 1.032, t=7.375, *P*=0.0018) (Fig. 6A and B). To identify the role of WT1 in regulating its downstream target gene TGF-β1, the expression of TGF-β1 was observed upon transfecting with si-WT1, and the transfection efficiency of si-WT1 was demonstrated by the relative expression of WT1(Supplimentary Figure 1A, B). These experiments revealed that, although WT1 expression in the si-WT1-transfected group was downregulated compared with the control group (t=3.085, P=0.037) (Fig. 6C and D), the expression of TGF-β1 was increased significantly (t=15.80, P<0.0001) (Fig. 6E and F). As compared with the si-NC group, the expression of CD11b was upregulated in the si-WT1 group (1.94±0.46 vs 18.36±0.12, t=59.96, *P*<0.0001) (Fig. 6G, H); in addition, the expression of CD14 was also upregulated in the si-WT1 group (3.29±0.32 vs 17.34±0.71, t=31.42, *P*<0.0001) (Fig. 6I, J). Taken together, these results suggested that WT1 can negatively regulate TGF-β1 expression of its downstream target gene TGF-β1, and that downregulation of WT1 expression by siWT1 resulted in the upregulation of TGF-β1 expression.

**Fig.6.**
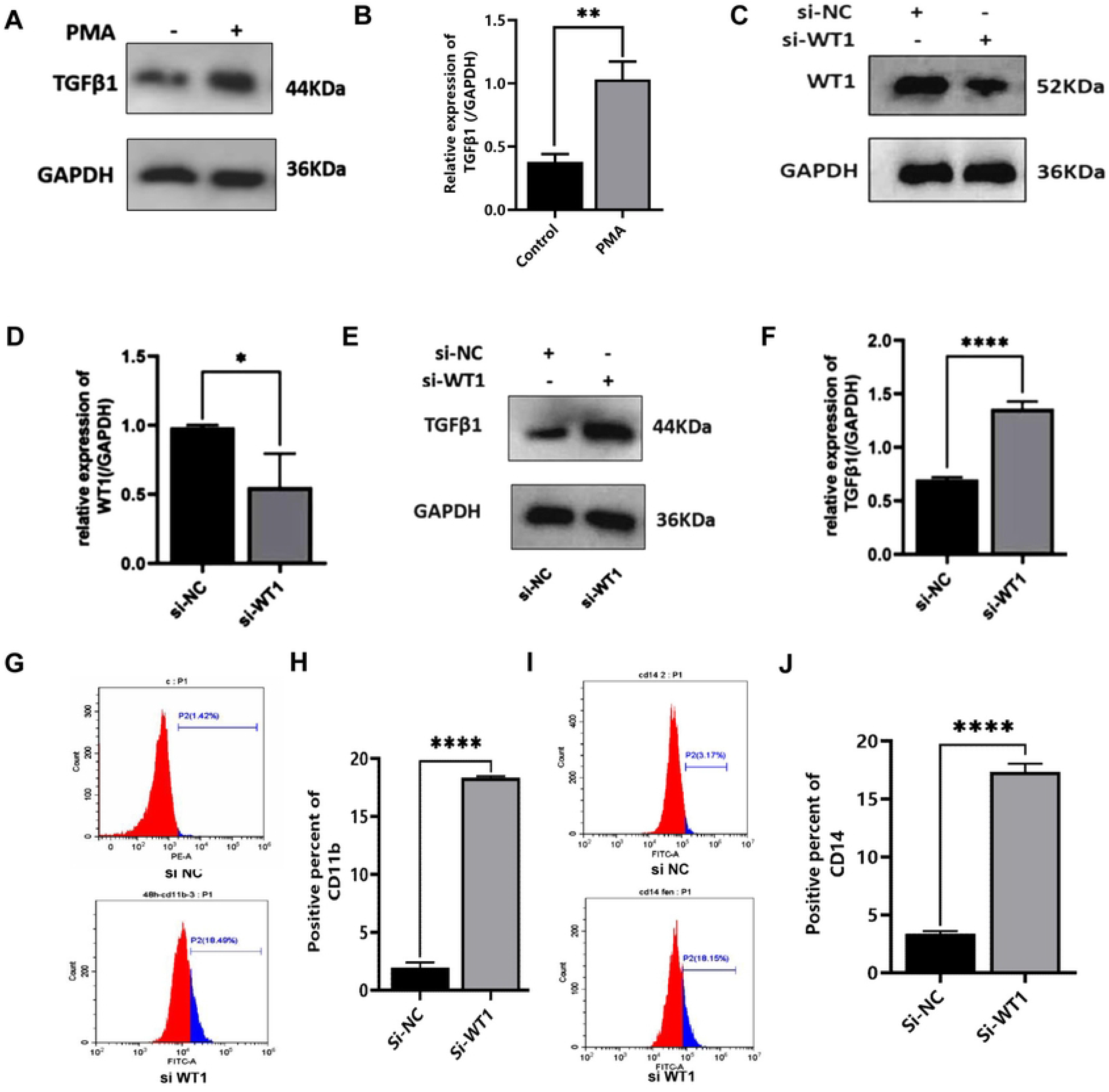
Targeting regulation of TGF-β1 expression by WT1 on committed differentiation of monocytes/macrophages. WT1 protein expression was detected by Western blotting assay, and the expression level of CD11b and CD14 differentiation antigen were detected by flow cytometry assay. A. Western blot was used to detect the protein level of TGF-β1 in THP-1 cells after PMA treatment; B. Changes in the mean level of TGF-β1 protein in THP-1 cells after PMA induction; C. Effect of si-WT1 on the expression of WT1 protein in THP-1 cells; D. Mean WT1 protein expression level in THP-1 cells transfected with si-WT1; E. Effect of si-WT1 on the expression of TGF-β1 protein in THP-1 cells; F. Mean expression level of TGF-β1 protein in THP-1 cells transfected with si-WT1.G.Expression of CD11b antigens by flow cytometry; H. Mean expression levels of CD11b differentiation antigens; I.Expression of CD14 antigens by flow cytometry; J.Mean expression levels of CD14 differentiation antigens. (* *P* < 0.05, *** *P* < 0.001, **** *P* < 0.0001)

### ChIP assay to detect WT1-targeted binding and regulation of TGF-β1 expression

To further verify whether WT1 targets the binding of TGF-β1, and regulates TGF-β1 expression, a ChIP assay experiment was designed. First, prediction analysis of the TGF-β1 promoter region for WT1 binding sites was performed, and a putative binding site in the region from -180 to +1 bp was identified, as has been mentioned above (Fig. 7A). Through performing database analyses and comparing the literature, the PCR amplification primer sequence of TGF-β1 was first determined (Fig. 7B). Four groups were set up in this experiment, including the PMA + IgG group (negative control group), PMA + input group (positive control group), Control + anti-WT1 group, and PMA + anti-WT1 group. The ChIP-PCR experiments yielded a negative value for the PMA+ anti-WT1 group, whereas that for the Control + anti-WT1 group was positive (Fig. 7C). Taken together, these results confirmed that WT1 can target both the binding and regulation of TGF-β1 expression.

**Fig.7.**
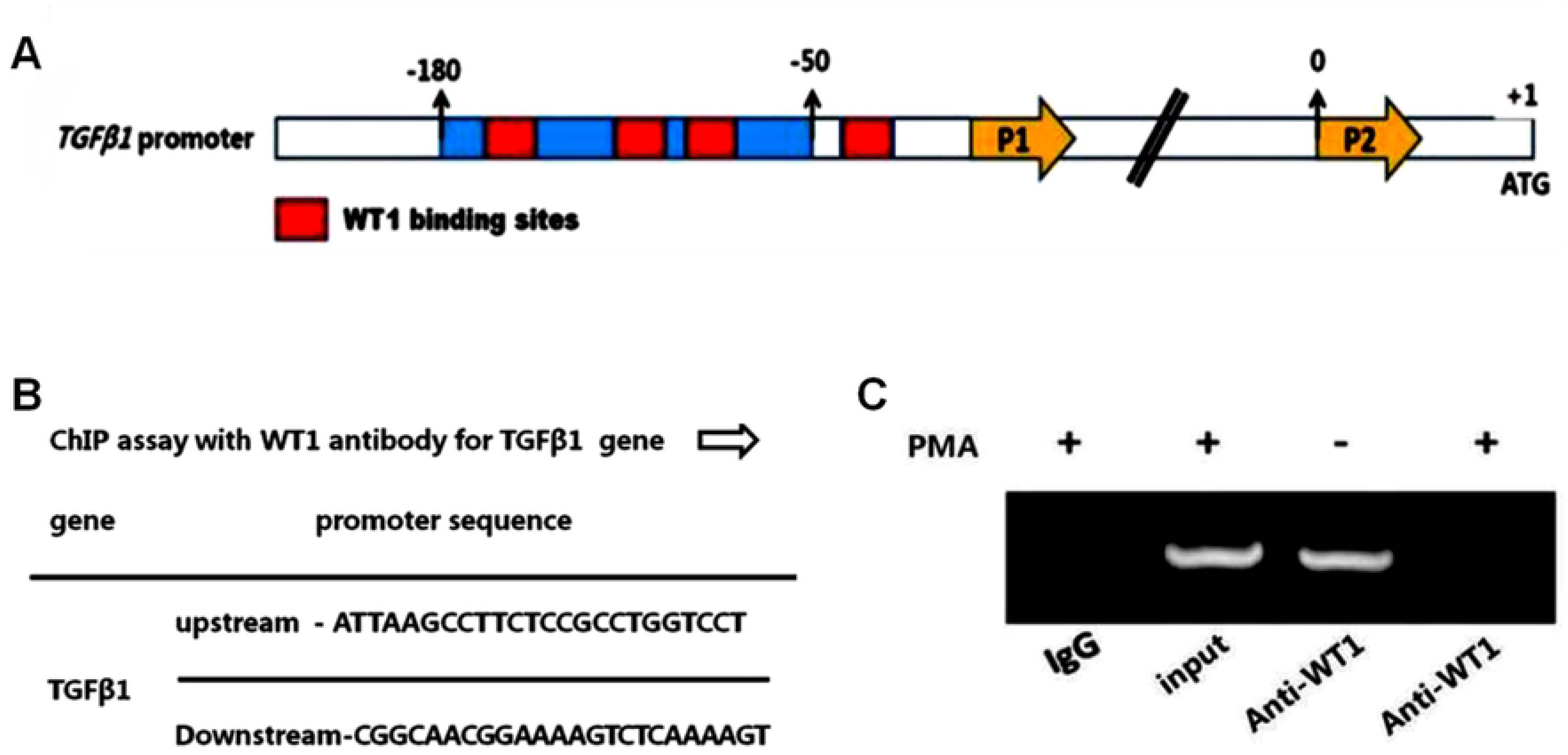
Chromatin immunoprecipitation (ChIP) assay for WT1 targeting and regulation of TGF-β1 expression To further verify whether WT1 targets binding and regulates expression of TGF-β1, Chromosome immunoprecipitation (ChIP) assay for targeting regulation of TGF-β1 by WT1. A. Prediction of the binding region of WT1-targeted TGF-β1 promoter; B. Primer sequence design for ChIP assay of TGF-β1; C. ChIP assay results of WT1 targeting TGF-β1.

Rescue experiment to investigate the role of miRNA-132-3p in inducing THP-1 cell differentiation via modulating the expression of TGF-β1. Subsequently, experiments were designed comprising the following experimental groups, namely the mimics-NC group, miRNA-132-3p mimics group and miRNA-132-3p mimics+si-TGF-β1 group. Western blotting assay was used to detect the protein expression of TGF-β1, and the transfection efficiency of si-TGF-β1 was demonstrated by the relative expression of WT1(Supplimentary Figure 1C, D). The results of western blot assay demonstrated that TGF-β1 expression in the miRNA-132-3p mimics-transfected group was higher compared with that of the mimics NC group (1.11±0.01 vs 0.59±0.12, t=8.177, P=0.0012). By contrast, the expression of TGF-β1 in the miRNA-132-3p mimics+si-TGF-β1 transfected group was lower compared with that in the miRNA-132-3p mimics only group (0.529±0.241 vs 1.108±, t=4.157, P=0.014) (Fig. 8A and B). Subsequently, the differentiation antigens of CD11b and CD14 were detected by flow cytometry assay. These experiments revealed that the expression of both CD11b and CD14 in the miRNA-132-3p mimics group was upregulated compared with that in the mimics NC group (t=41.23, P<0.0001; and t=22.96, P<0.0001 for CD11b and CD14, respectively). In addition, the expression of the CD11b antigens in the miRNA-132-3p mimics + si-TGF-β1-transfected group was downregulated compared with that in the miRNA-132-3p mimics group (Fig. 8C and D), and the differences were found to be statistically significant (20.04±0.66 vs 8.49±0.95, t=17.18, P<0.0001). The expression of the CD14 antigens in the miRNA-132-3p mimics + si-TGF-β1-transfected group was downregulated compared with that in the miRNA-132-3p mimics group (Fig. 8E and F), and the differences were found to be statistically significant (24.31±1.28 vs 9.13±2.50, t=9.327, P=0.0007, respectively). Taken together, these results indicated that miRNA-132-3p is able to positively regulate TGF-β1 expression, and an elevated level of miRNA-132-3p expression upregulates TGF-β1 expression, thereby promoting THP-1 cell-directed macrophage differentiation, whereas siTGF-β1 causes a reversal of this role of miRNA-132-3p in modulating expression of CD11b and CD14 antigens.

**Fig.8.**
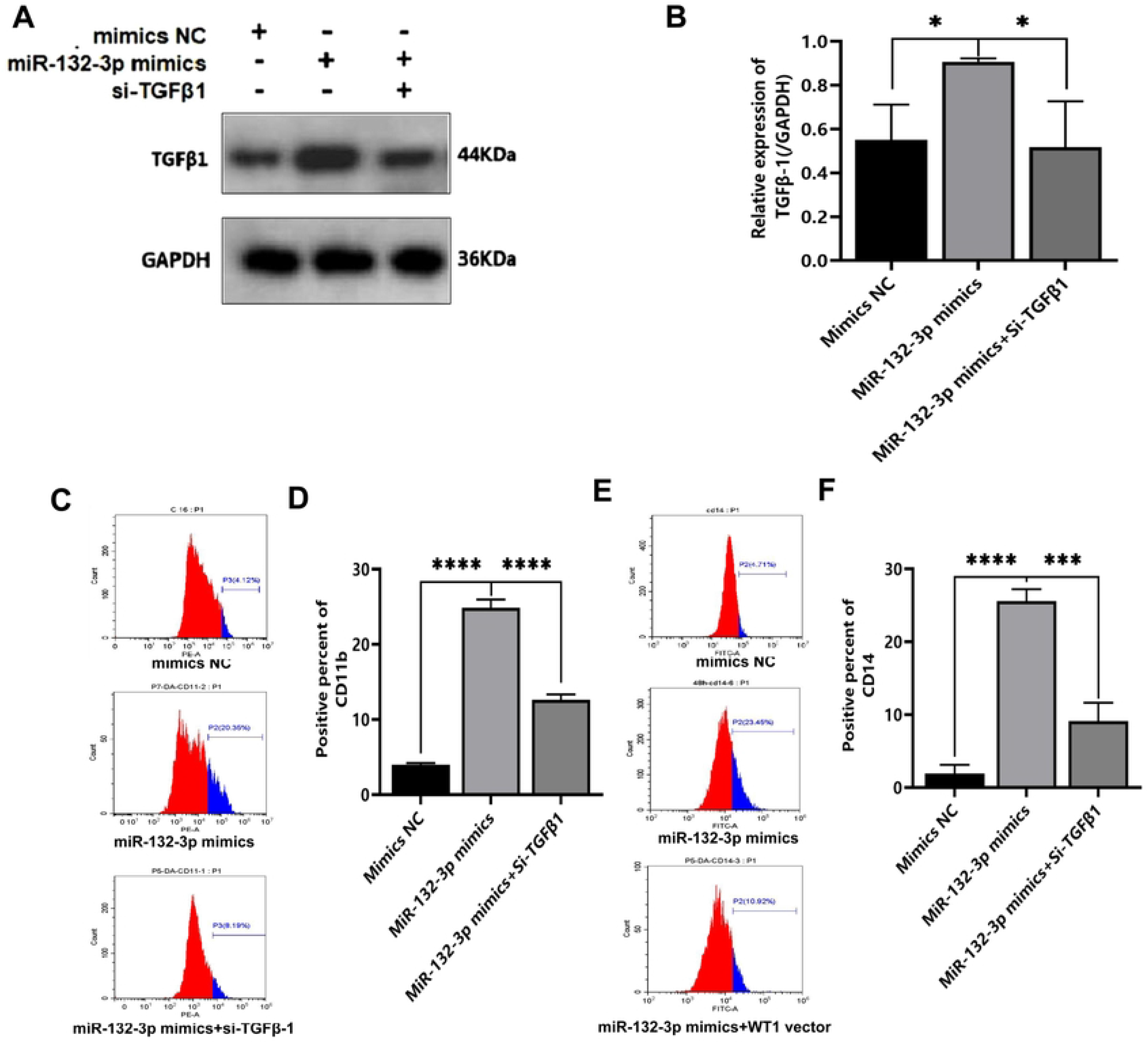
Rescue assay results of miRNA-132-3p targeting TGF-β1 to induce differentiation of THP-1 cells miRNA-132-3p mimics-NC group, miRNA-132-3p mimics group, and miRNA-132-3p mimics+si-TGF-β1 group three groups were used to try rescue assay about miRNA-132-3p modulating expression of TGF-β1. Western blotting assay was used for detecting protein expression of TGF-β1. A. Results of Western blot assay for TGF-β1; B. Changes in the mean expression level of TGF-β1 protein; C. The CD11b expression by flow cytometry; D. Mean expression level of CD11b differentiation antigen; E. The CD14 expression by flow cytometry; F. Mean expression levels of CD14 differentiation antigens. (* *P* <0.05, ** *P* <0.01)

The miRNA-132-3p/WT1/TGF-β1 signaling pathway is involved in inducing differentiation of THP-1 cells towards macrophages induced by PMA. Considering all the results obtained thus far, it was possible to conclude that PMA can induce the committed differentiation of THP-1 cells towards macrophages. During this committed differentiation process, miRNA-132-3p expression is upregulated, WT1 expression is reduced, and TGF-β1 expression is also upregulated. Moreover, these three genes have been demonstrated to be involved in this committed macrophage differentiation of leukemia cells. Through target gene prediction and the dual-luciferase reporter gene assay experiments, it was confirmed that miRNA-132-3p regulates the level of promoter activity of the WT1 gene through targeting its 3’-UTR region, thereby regulating the committed macrophage differentiation of leukemia cells. Subsequently, it was also demonstrated that miRNA-132-3p can regulate TGF-β1 by targeting WT1, thereby promoting the differentiation of THP-1 cells towards macrophages. In conclusion, the miRNA-132-3p/WT1/TGF-β1 axis has been shown to contribute to the polarization of THP-1 cells towards macrophages (Fig. 9).

**Fig.9.**
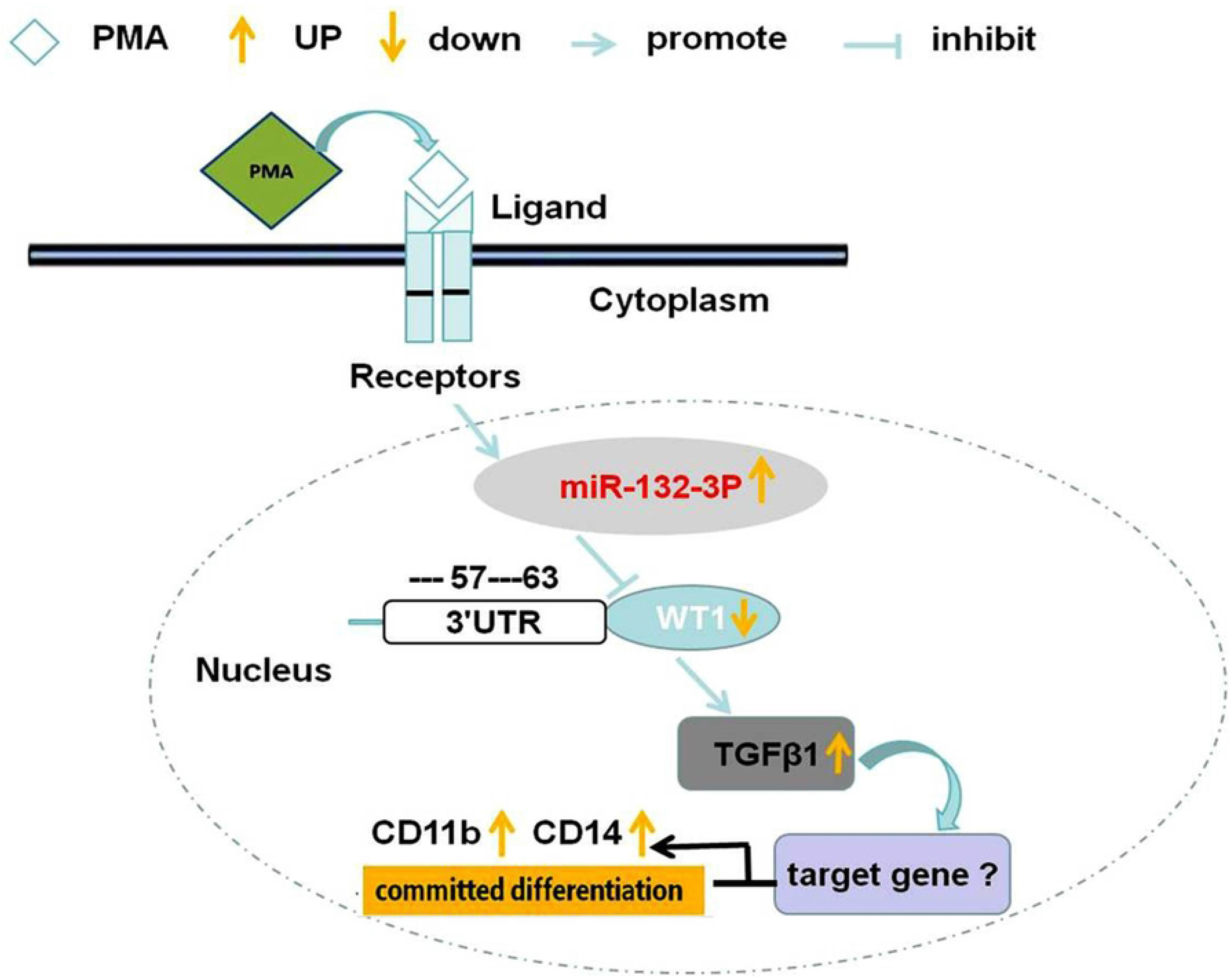
Schematic of the signaling pathway of the miRNA-132-3p/WT1/TGF-β1 axis contributed to inducing differentiation of THP-1 cells into macrophages. In THP-1 cells, PMA could induce the upregulation of miRNA-132-3p, the upregulation of miRNA-132-3p results in the downregulation of WT1 by sponge action with binding on its 3’UTR, the down regulation of WT1 contributes to the upregulation of TGF-β1, then the polarization of macrophages is induced, which is indicated by the upregulation of CD11b and CD14 expression. Therefore, the miRNA-132-3p/WT1/TGF-β1 axis is involved in the committed differentiation of THP-1 leukemia cells into macrophages.

## Discussion

AML is a malignant clonal disease of hematopoietic stem cells that blocks the differentiation of myeloid cells (46,47). Previously, the treatment of leukemia has mainly relied on drug-targeted therapy and chemotherapy. Although some progress has been made clinically in terms of the use of therapeutic drugs in the early stage, with the continuous use of these drugs, it is inevitable that drug resistance, recurrence, and clear toxic side effects will occur. Therefore, there is an urgent need to explore other safer and more effective treatment methods, and to develop novel drugs. Interestingly, it has been demonstrated that induction differentiation therapy is an ideal therapeutic method for the treatment of leukemia, and has been successfully used in the clinic for the treatment of patients with APL (48-52). However, to date, leukemia cells have only been successfully induced to differentiate into granulocytes, and no significant breakthroughs have been achieved in inducing differentiation towards monocytes and macrophages. This is mainly accounted for by the fact that the underlying molecular mechanism that determines the directed differentiation of macrophages remains poorly understood.

The quest to unravel the underlying molecular mechanisms of leukemia differentiation disorders is of crucial importance, as it provides an important basis for finding targeted therapeutic targets. However, the selection of the research entry point is directly related to whether the key signaling pathways of leukemia cell differentiation disorders can be uncovered, especially the discovery of key regulatory molecules or important signaling pathways that are associated with the differentiation of the leukemia cells. According to previous studies, certain oncogenes or tumor suppressor genes act as important regulatory factors for the abnormal proliferation or differentiation of tumor or leukemia cells. WT1, as an effective transcription regulatory factor, has been shown to be closely associated with the proliferation, apoptosis, differentiation and survival of normal or tumor cells (53-55). In clinical studies of patients with AML, a previous study suggested that WT1 is an important factor associated with the occurrence and development of leukemia. (56). WT1 has been found to harbor mutations and to be expressed to differing degrees in different subgroups of AML, and consequently, it is used as a biomarker for the diagnosis of AML, monitoring of MRD, clinical management, and molecular remission or recurrence detection. Another study (21) has explored its application as a potential biological target for immunotherapy. An abnormal expression of WT1 has also been observed in leukemia mother cells of B- or T-lineage acute lymphoblastic leukemia (ALL), AML, critically ill patients with chronic myeloid leukemia, and patients with myelodysplastic syndrome (20). In addition, the majority of recently published studies [e.g., (22)] have suggested that wild-type WT1 protein is highly expressed in acute myeloid leukemia, even in patients with WT1 mutations. Therefore, WT1 has been shown to have opposite roles in leukemia, namely being an oncogene when aberrantly expressed, and a tumor suppressor gene when mutated. To date, the exact role of WT1 in the development of leukemia remains unclear, or at least has only been partly elucidated, due to the complexity of WT1 regulation, given that WT1 has a variety of different roles, serving as both an oncogene and a tumor suppressor. The complexity of its functions means that a lot of further research studies are required, approaching the study of WT1 from a range of different aspects (25). On the other hand, the high expression status of WT1 already provides a useful opportunity for the development of targeted immunotherapy and other therapies. Targeted therapy for WT1 mainly involves the use of wild-type or modified major histocompatibility complex (MHC) class I restrictive WT1 peptides to produce vaccines against WT1-overexpressing tumor cells and leukemia mother cells (23). In addition, another study (24) has reported on the development of monoclonal antibodies targeting the WT1/MHC-I peptide complex expressed on the surface of leukemia mother cells. Therefore, although WT1 is well established as one of the oncogenes closely associated with tumors or leukemia, to date, few studies have reported on its important role in the targeted differentiation of leukemia. Due to its involvement in the proliferation and differentiation processes of tumor cells, however, it may be speculated that it exerts its role through participating in a certain signaling pathway.

Accordingly, we hypothesized that WT1 may be involved in the directed differentiation of leukemia cells. In the preliminary experiments, differential gene expression between THP-1 cells and normal peripheral blood monocytes was analyzed through RNA-seq. The results obtained confirmed that WT1 is abnormally overexpressed in leukemia cells, and enrichment analysis revealed that it is associated with the process of cell differentiation. In order to further reveal the role of WT1 in the directed macrophage differentiation of leukemia cells, in the present study, THP-1 cells were induced by PMA to establish a representative cell model for macrophage-committed differentiation, as described previously (53,54). It was found that PMA could induce the committed differentiation of THP-1 leukemia cells towards macrophages, as evaluated by the upregulation of CD11b and CD14 antigens. In our investigated model of cell differentiation into macrophages induced by PMA, we further found that the expression of WT1 protein was gradually decreased, accompanied by a significant increase in the expression of CD11b and CD14 differentiation antigens in THP-1 cells. After transfecting the cells with siWT1 to interfere with WT1 expression, these two differentiated antigens were significantly increased. These results preliminarily suggested that WT1 is involved in the committed differentiation of the THP-1 cells; however, further in-depth research was required to determine how WT1 participates in the ‘switch’ from THP-1 cells into macrophages. Therefore, the present study has shown that WT1 inhibitors or siWT1 may be therapeutic agents to potentially treat patients with leukemia by inducing the committed differentiation to macrophages.

miRNAs are endogenous noncoding small RNAs that usually target the 3’-UTRs of specific target mRNAs to regulate the expression of target genes (57), resulting in the regulation of cell differentiation or apoptosis (58). A number of different studies have demonstrated that certain miRNAs may function as potential biomarkers for cancer diagnosis and treatment (59). Usually, miRNAs exert their role through a ‘sponging’ action to modulate downstream genes. This serves the purpose of regulating the expression of downstream oncogenes or tumor suppressors, such as targeting WT1 to regulate the proliferation of tumor cells (60,61). To reveal the specific role of WT1 in the committed differentiation, in the present study, its upstream microRNAs were predicted and validated. Based on the prediction results that were obtained, it was possible to surmise that, in the process of PMA inducing THP-1 cells to polarize into macrophages, miRNAs may exert a role through the targeted regulation of WT1. To confirm this hypothesis, RNA-seq was used as the technology to screen and analyze differentially expressed miRNAs both before and after PMA induction. On the other hand, based on the fact that miRNAs often exhibit a negative correlation with the expression levels of downstream target gene mRNAs, we focused on analyzing the differentially expressed miRNAs that were upregulated following exposure to PMA. In addition, considering the complementary binding characteristics of miRNAs with specific downstream target gene promoters, potential miRNAs that could bind to WT1 were predicted using the Targetscan and mirDB databases. Furthermore, literature research results on the targeted regulation of WT1 were manually searched using the PubMed/MEDLINE database. After performing an intersection analysis of these three related investigations, together with performing an additional RT-qPCR analysis for identification purposes, miRNA-132-3p emerged as the miRNA most likely to regulate WT1 expression. Therefore, miRNA-132-3p was identified as the potential upstream miRNA targeting WT1, which was predicted using the Targetscan database. Dual luciferase reporter gene assay further confirmed that WT1 is truly a downstream gene of miRNA-132-3p, and the application of miRNA-132-3p mimics resulted in a decrease in WT1 expression, accompanied by an upregulation of the CD11b and CD14 antigens. After simultaneously transfecting the cells with miRNA-132-3p mimics and WT1 vector, the expression level of WT1 was effectively restored, antagonizing the effect of the miRNA-132-3p mimics, manifested by a significant downregulation of the CD11b and CD14 differentiation antigens. Taken together, these results suggested that miRNA-132-3p is involved in the PMA-induced macrophage-directed differentiation of THP-1 cells through regulating the expression level of WT1. Moreover, the results showed that miRNA-132-3p may be a potential tumor-suppressive miRNA that functions similarly to other tumor-suppressive miRNAs in blocking and alleviating the progression of leukemia (62).

After having confirmed the identity of the upstream miRNA of WT1, subsequently, whether WT1 has a role in targeting downstream target genes became the focus of our research. Based on a literature analysis, as an important member of the TGF family, TGF-β1 was identified not only as a participant in cell proliferation, differentiation, and apoptosis, but it also suppresses the proliferation of tumor cells, or induces differentiation of leukemia cells (63,64). Therefore, we hypothesized that WT1 may regulate the differentiation of THP-1 cells into macrophages via regulating the TGFβ/SMAD pathway. Experiments were designed to investigate whether WT1, as a transcription factor, binds the promoter of TGF-β1 to modulate the differentiation of THP-1 cells. That WT1 may be a possible transcription factor targeting the promoter of TGF-β1 was first identified through database prediction analysis. Subsequently, a ChIP assay was performed to establish whether WT1 could interact with the promoter of TGF-β1, and whether WT1 could negatively regulate the expression of TGF-β1, as a target gene, to promote the differentiation of THP-1 cells into macrophages. Furthermore, we sought to investigate whether miRNA-132-3p could affect the differentiation of THP-1 cells via regulating TGF-β1 expression. Taken together, the results of these experiments indicated that miRNA-132-3p mimics were able to positively regulate TGF-β1 expression, and the interference (or rescue) experiment, wherein we investigated the effect of transfection with siTGF-β1 on miRNA-132-3p mimics in terms of regulating committed differentiation, further demonstrated that miRNA-132-3p indeed exerts its role through regulating the expression of TGF-β1.

Although it has been preliminarily shown that miRNA-132-3p, WT1 and TGF-β1 have important roles in the PMA-induced differentiation of THP-1 cells into macrophages by way of upstream or downstream regulation, further studies are required in order to fully delineate the underlying mechanism, and to reveal the significance of their regulatory associations. For example, the upstream genes or transcription regulatory proteins that regulate the expression of miRNA-132-3p need to be identified, and the manner in which TGF-β1 regulates its downstream target genes (and its complete role in performing this function) still needs to be explored. Secondly, given that it is typically encountered as the downstream gene of the TGF-β1 signaling pathway (65,66), the role of the Smad family in inducing differentiation of leukemia cells into macrophages needs to be elucidated. In addition, in order to clarify the changes observed in these three genes, and the significance of the signaling axis that they regulate, the required in vivo animal studies and the detection of primary leukemia cells in clinical patients will be important aspects of our subsequent experiments. In spite of this, the present work, which was mainly focused on THP-1 cells, did achieve the hoped-for results. Furthermore, to corroborate the work done with the THP-1 cells, the K562 or U937 cell line will be utilized in future studies to investigate whether the miR-132-3p/WT1/TGF-β1 signaling axis exerts a similar role in the committed differentiation into macrophages.

In conclusion, the current research results indicated that WT1 is highly expressed in malignant cells of patients with hematological malignancies, or a significant proportion of patients carry inactivated mutations of this gene (67). For example, abnormal expression of WT1 was observed in leukemia cells of patients with B or T lineage acute lymphoblastic leukemia (ALL), acute myeloid leukemia (AML), chronic myeloid leukemia (CML), and myelodysplastic syndrome (MDS) (68). However, the exact role of WT1 in the development of leukemia is still not fully understood, and WT1 sometimes has two contradictory roles in leukemia. It is an oncogene when it is abnormally expressed, a tumor suppressor gene when it mutates, and is related to the specific tissue environment. The complexity of its functions, multiple isomers, and the lack of ideal research models all pose challenges to further clarify its functional characteristics (69,70). So far, many studies on the mechanism of WT1 have focused on its function as a transcription factor and the role of its many isomers. However, recent work from AML large-scale genome research has revealed a new role of WT1 in epigenetic regulation (71). On the other hand, just due to the abnormal expression or isomer of WT1, some scholars have confirmed that it can serve as an indicator for minimal residual disease (MRD) monitoring and prognosis judgment (72). The present study has demonstrated, to the best of our knowledge for the first time, that the miRNA-132-3p/WT1/TGF-β1 signaling axis is an important pathway that is involved in the committed differentiation of THP-1 leukemia cells into macrophages, providing a good theoretical basis underpinning future in-depth research on the clinical significance of this signaling axis. The importance of the miRNA-132-3p/WT1/TGF-β1 signaling axis as a prognostic indicator for patients with leukemia needs to be validated through further clinical studies, and the potential value of targeted intervention therapy needs to be validated at the animal level first. Inducers or inhibitors of this signaling pathway also have the potential to become targeted intervention drugs.

## Acknowledgements

Not applicable.

## Availability of data and materials

The datasets used and/or analyzed during the current study are available from the corresponding author on reasonable request.

## Authors’ contributions

KHB, GSJ, QL, CZW, and ZMW made substantial contributions to the conception and design and also critically reviewed the study, ZMW, CZW, DFZ, XDW, QL, YHW, RJS and XLS performed the experiments. ZMW, CZW, YHW and XLS analyzed the data and wrote the manuscript. KHB and GSJ confirm the authenticity of all the raw data. All authors read and approved the final manuscript.

## Ethics approval and consent to participate

Not applicable.

## Patient consent for publication

Not applicable.

---

Supplementary Figure 1

The transfection efficiency of si-WT1, si-TGF-β1 and WT1 vector detected by the relative expression of their proteins. A. The WT1 expression by Western blotting assay in THP-1 cells transfected with siNC and si-WT1; B. The mean relative expression of WT1 in THP-1 cells transfected with siNC and si-WT1; C. The TGF-β1 expression by Western blotting assay in THP-1 cells transfected with siNC and si-TGF-β1; D. The mean relative expression of TGF-β1 in THP-1 cells transfected with siNC and si-TGF-β1; E. The WT1 expression by Western blotting assay in THP-1 cells transfected with empty vector and WT1 vector; F. The mean relative expression of WT1 in THP-1 cells transfected with empty vector and WT1 vector.

